# DNA binding drives phase separation of the Gcn4 bZIP domain and reveals its conformational ensemble in the diluted and condensate phases

**DOI:** 10.64898/2026.02.10.704833

**Authors:** Antonino Calio’, Florian Turbant, Mark D. Tully, Satyan Sharma, Gioacchino Schifino, Antonietta Parracino, Judith Peters, Annalisa Pastore

**Affiliations:** Université Grenoble Alpes, CNRS, LiPhy, 38400 Grenoble, France; European Synchrotron Radiation Facility, 71 Avenue des Martyrs, Grenoble, France; Department of Chemistry-BMC, Uppsala University, Uppsala, S-751 23 Sweden; Institut Laue Langevin, 38042 Grenoble, France; Institut Universitaire de France, 75231 Paris, France; King’s College London, London, UK; Elettra Sincrotrone Trieste, Basovizza, Italy

**Keywords:** ensemble optimization method, intrinsically unfolded proteins, SAXS, synchrotron radiations, transcription factors

## Abstract

Liquid-liquid phase separation is widely invoked in transcriptional regulation, yet prevailing models attribute condensate formation primarily to intrinsically disordered activation domains rather than structured DNA-binding motifs. Here, we overturn this view by demonstrating that the isolated basic leucine zipper (bZIP) domain of the yeast transcription factor Gcn4 undergoes robust DNA-induced phase separation in the complete absence of its activation domain. Using small-angle X-ray scattering in combination with all-atom molecular dynamics simulations and ensemble optimization, we directly resolve the conformational landscape of the Gcn4 bZIP–DNA complex across coexisting dilute and condensed phases. Beyond the canonical uninterrupted helical conformation captured in crystal structures, we identify a previously unrecognized minor population featuring a pronounced helical kink at the basic region–leucine zipper junction. These findings establish DNA binding as a sufficient physical driver of bZIP phase separation and demonstrate that small-angle scattering can quantitatively interrogate protein conformational ensembles within biomolecular condensates, opening new avenues for the structural chemistry of phase-separated systems.

## Introduction

It is now well established that cells are not merely bags of freely floating molecules or rigid membrane-surrounded compartments but contain a rich interior of dynamic and organized environments. Some of them are not defined by membranes but through a process known as liquid-liquid phase separation (LLPS) (Alberti and Dormann, 2019). This process is somewhat analogous to the way oil droplets separate from water. The components of the liquid droplets (or condensates), that mostly comprise proteins and nucleic acids, can move around within them, exchange with the outside environment, and fuse with other droplets, usually in a reversible fashion in response to changes in cellular conditions. This makes them ideal for regulating processes like gene expression, RNA metabolism, and stress responses (Chen et al., 2026; He at al., 2025). However, while LLPS is normally a functional and healthy process, under the wrong conditions, such as stress, aging, or mutation, proteins can lose their ability to remain dynamic and soluble; and harden into solid or gel-like aggregates that are usually irreversible. As a result, they may interfere with normal function sequestering essential proteins and making them unavailable to fulfil their functions. This aggregation is a hallmark of many neurodegenerative diseases, such as Alzheimer’s disease, Huntington chorea and amyotrophic lateral sclerosis (Andrade et al., 2025; Dobson et al., 2020).

Transcription activation, the first and essential step in gene expression, is one of the processes that would strongly benefit from LLPS formation. It is in fact clear that dynamic crowding and confinement would greatly help to increase the local concentration of transcription factors and coactivators at target genes (Chen et al., 2026; He at al., 2025). This molecular clustering would drastically increase the efficiency with which RNA polymerase II is recruited to promoters and enhancers, thereby stimulating transcription initiation. LLPS would also provide a way to integrate regulatory signals. Because condensates form only when protein or RNA concentration reaches a threshold, they could act as natural molecular switches. However, while these are reasonable assumptions, whether transcription is mediated by biomolecular condensates or by non-stoichiometric soluble assemblies distinct from phase separated condensates and whether LLPS is necessary, sufficient, or consequential for transcription is still intensely debate (Stortz et al., 2024; Narlikar et al., 2021; McSwiggen et al., 2019). Recently, a study presented strong evidence that the two processes could be mutually compatible. The authors showed that the transcriptional activities of several variants of the yeast protein Gcn4 are explained both by their abilities to phase separate with the coactivator Med15 of the Mediator complex and by their abilities to form soluble complexes (Bremer et al., 2025). This study has brought Gcn4 back under the spotlight, a protein of 281 amino acids that was discovered in the early 90s (Lucchini et al. 1984) and used ever since for completely different purposes.

Gcn4 contains, in addition to long intrinsically disordered regions that span the transcription activation domain, a prototypical example of a coiled-coil motif which was renamed *ad hoc* leucine zipper (Agre et al., 1989). This motif is a dimerization domain formed by two parallel amphipathic α-helices containing heptad repeats with leucines in the first and fourth positions, which zip together via hydrophobic interactions to form a coiled-coil dimer (O’Shea et al., 1991; Saudek et al., 1991; Ellenberger et al. 1992). The C-terminal leucine zipper of Gcn4 is preceded by a basic region that binds DNA with high specificity forming a bZIP motif. The basic region is intrinsically disordered although with a considerable helical tendency but forms an uninterrupted helix as soon as DNA binds (Weiss et al., 1990; Saudek et al., 1990; Saudek et al., 1991). Gcn4 bZIP inspired new research in the field of protein design because, when the rules to form leucine zippers were clarified, the leucine zipper motif was widely used to build dimeric, trimeric and tetrameric coiled-coils (O’Shea et al., 1993; Harbury et al., 1993; Harbury et al., 1994). A detailed description of the dynamics of Gcn4 bZIP was also obtained through the combination of extensive molecular dynamics simulations and relaxation studies by nuclear magnetic resonance (Bracken et al., 1999; Gill et al., 2006; Robustelli et al., 2013). This knowledge has made the motif ideal for designing synthetic transcription factors, stabilizing fusion adaptors or modular protein interaction domains (Weissenhorn et al., 1997; Wolfe et al., 2003; Deiss et al., 2014). Many human transcription activators contain bZIP motifs, making Gcn4 an excellent model also for higher organism transcription.

While working on Gcn4 bZIP with the aim of characterizing its interaction with plasmids with different levels of supercoiling, we discovered that the isolated bZIP domain undergoes LLPS in the presence of DNA. This was intriguing because Bremer et al. (2025) had reported phase separation only for constructs containing the transcription activator domain. We then set up to investigate the phenomenon. To better understand the nature of this phase transition and characterize the structure of the Gcn4 complex with DNA in solution and in the liquid droplets, we used state-of-the-art small angle X-ray scattering (SAXS) techniques to study the structure of the Gcn4 bZIP both in isolation and in the presence of its cognate DNA. For this purpose, we used the ARG-1A DNA aptamer, which has nanomolar affinity and specificity for Gcn4 (Coey and Clark, 2022). We demonstrated that, thanks to the advanced techniques used, not only can we accurately describe the conformational changes occurring when the Gcn4 bZIP dimer binds to DNA according to already acquired knowledge, but we can also provide novel quantitative information about the conformations of the protein in the complex (and thus about its dynamics), both in solution and in the condensate droplets. Our results show how sensitive SAXS methods are to describe molecular structure and dynamics also in molecular condensates and provide brand new information about the structure of Gcn4 bZIP in different solution states.

## Materials and Methods

### Protein production

The plasmid pGCNK58 for *E. coli* expression of a construct of Gcn4 containing both the basic region and the leucine zipper domain (bZIP) without any tag was kindly provided by Prof. A. Palmer (Gill et al., 2016). The protein was expressed and purified according to previously well-described protocols (Gill et al., 2016). In short, the protein was expressed in *E. coli* BL21 (DE3) cells, transformed and grown in 4 L of Luria Broth media at 37 °C until reaching an optical density (OD_600_) of 0.5. Protein expression was induced with 1 mM IPTG and allowed to proceed for approximately 4 h. Lysis was achieved by sonication in HEPES 25 mM, 50 mM NaCl pH 7.5. The protein was purified by Ion Exchange Chromatography on AKTA Go system with Cityva HiTrap SP FF 5 ml column. Elution was performed by NaCl gradient from 50 mM to 1 M. Fractions were checked by SDS-PAGE, pooled, concentrated to 1.2 mM, and dialyzed in 25 mM HEPES and 50 mM NaCl. An attempt was also carried out by adding a further step of purification by reverse phase HPLC according to the available protocol. Purity was checked by SDS-PAGE (**Figure S1 of Suppl. Mat**.). Since the final purity was estimated to be >98% before the reverse-phase chromatography, this step was not used for the main production batch. Protein concentration, determined by UV spectroscopy, was defined with respect to the monomer. Conclusive identification of the protein and confirmation of the correct folding were obtained by MALDI TOF Mass Spectrometry (using the advanced platform of the Institut de Biologie Structurale of the EPN campus in Grenoble) and by circular dichroism (**Figures S2 and S3 of Suppl. Mat.)**. The ARG-1A aptamer (5’ TTTTTAGTGACTCATGTCGCATTT 3’ in which the RNA-binding motif is underlined) was purchased from Sigma Aldrich (St Luis, MI, batch HA 16993262).

### Small Angle Scattering measurements

Small-angle X-ray scattering (SAXS) experiments were carried out at the BM29 beamline of the European Synchrotron Radiation Facility (ESRF, Grenoble, France, https://doi.org/10.1107/S1600577522011286). Data were collected on freshly purified Gcn4 leucine zipper dimer (0.75 mM or 1.5 mM monomer concentration) and on the complex between Gcn4 and the synthetic double-stranded DNA aptamer ARG-1A (0.62 mM). All samples were prepared in a measurement buffer containing 25 mM MES pH 5.75, 150 mM NaCl, and 2.5 mM MgCl_2_. Buffer scattering was recorded under identical conditions and subtracted from the sample scattering profiles. The beamline was operated at a wavelength of 0.99 Å (12.5 keV), with a sample-to-detector distance of 2812 mm on a PILATUS3 2M detector, covering a scattering vector range of 0.007-0.55 Å^-1^, where q = 4π sin(θ)/λ. Ten successive frames of 1 s exposure each were acquired for every sample in batch mode at 20 °C, with beam current at 200 mA. Repeated buffer measurements were taken before and after each sample and subtracted from the sample scattering to remove background contributions. Absolute scaling of the forward scattering I(0) was performed by water calibration. To assess potential concentration-dependent effects, scattering data were collected at multiple protein concentrations when possible. Exposure times were optimized to minimize radiation damage, and data were systematically checked for radiation induced aggregation by comparing sequential frames. The resulting scattering curves were averaged, buffer subtracted, and further analyzed using FreeSAS, BioXTAS RAW, and ATSAS software packages (Tully et al., 2021; Hopkins et al. 2024; Manalastas-Cantos et al., 2021). The radius of gyration (Rg) and forward scattering intensity (I(0)) were determined by Guinier analysis, while the pair distance distribution function P(r) was determined using GNOM (Svergun 1992). Molecular weights in solution were estimated from I(0).

To compare experimental data with known structural information, theoretical scattering curves were computed from the crystal structure of Gcn4 bound to DNA (PDB ID: 1YSA) using CRYSOL (Svergun et al., 1995). Since Gcn4 undergoes a transition to a more disordered state in absence of DNA (Weiss et al., 1990), flexible refinement of the model was carried out using SREFLEX (Panjkovich and Svergun, 2016). Ab initio electron density reconstructions were performed with DENSS to validate model fitting (Grant, 2018)

### Molecular dynamics Simulations

Full details of these simulations are described elsewhere (Mattossovich et al., in preparation). In short, the structure of a complex of a linear DNA 336 bp long containing the ARG-1A sequence was built using the software NAB implemented in AmberTools22 using the crystal structure (PDB code 1YSA) as the template from which only the coordinates of the GCN4 bZIP and the ARG-1A sequences were extracted. The generated atomistic structures were subjected to two-step minimization, first using all atom coordinate restraints and second without restraints. Each minimization was done for 1000 steps using AMBER22 with the ff99SB force field with parambsc0 to describe DNA. All-atom simulations were performed with the GROMACS suite (Abraham et al., 2015). The solute was placed in a rectangular box. The minimum distance between any solute atom to the box edges was set to 4 nm. The system was subsequently solvated in TIP3 water (Jorgensen et al., 1983). The water molecules that were within 1.2 nm of the solute were retained to preserve accurate solute-solvent interactions while the remaining bulk water was replaced by the coarse-grained WT4 water model, following the multiscale protocol defined in the SIRAH force field (Machado et al., 2019). An ion concentration of 150 mM was imposed. This setup allows efficient multiscale molecular dynamics simulations, preserving atomistic details near the biomolecule while significantly reducing computational costs in the bulk solvent. The atomistic system used the AMBER99SB-ildn all-atom force field (Lindorff-Larsen et al., 2010), and included the parmbsc1 correction for DNA (Ivani et al., 2016). The solvated system was subjected to 5000 steps of steepest-descent minimization. Next, NVT equilibration was ran for 4 ns with positional restraints on solute. This was followed by NPT equilibration for 20 ns, with gradually weaker positional restraints only on the solute backbone atoms (1000 kJ/mol/nm^2^ for 2 ns, 100 kJ/mol/nm^2^ for 8 ns and 10 kJ/mol/nm^2^ for another 10 ns). The time step was 2 fs. Pressure was controlled at 1 bar via the Parrinello-Rahman barostat (Parrinello et al., 1981). Temperature was maintained at 300 K using the V-rescale thermostat (Bussi et al., 2007). Finally, production simulations were carried out for 100 ns. During the simulation, the trajectories were written every 10 ps and the coordinates of ARG-1A and Gcn4 bZIP were selected from each of the frames for SAXS analysis. The secondary structure elements were assigned using the DSSP (Dictionary of Secondary Structure for Proteins)-software (Kabsch and Sander, 1983). We estimated the flexibility of the protein structure by calculating the root-mean square fluctuation (RMSF). The secondary structure and RMSF analysis were done independently for each monomer of the dimeric Gcn4 bZIP. To quantify intra-monomer helix bending, we monitored an angle (θ) between two helix-axis vectors defined from centers of mass (COM) of the backbone atoms of selected dihedrals defined as follows: 235:C-238:C (N1), 245:C-248:C (C1), 253:C-256:C (N3), 272:C-275:C (C3) for the first monomer, and 235:D-238:D (N2), 245:D-248:D (C2), 253:D-256:D (N4), 272:D-275:D (C4) for the second monomer. The residues numbering follows the numbering used in the 1YSA PDB. The two angles were then calculated using 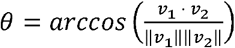, where *ν*_1_ is the vector from N1 to C1 and *ν*_2_ that from N3 to C3 for the first monomer; *ν*_1_ is the vector from N2 to C2 and *ν*_2_ from N4 to C4 for the second monomer.

### Estimate of the ensemble conformational heterogeneity

The Ensemble Optimization Method (EOM) was employed to analyze the conformational heterogeneity of the Gcn4 bZIP-DNA aptamer complex in solution, (Bernardó et al., 2007; Tria et al., 2015). The starting coordinates of the Gcn4 bZIP and ARG-1A complex were further refined and converted to PDB coordinates for subsequent analysis. Theoretical scattering curves were calculated for each conformer using FFMAKER (ATSAS suite), and the resulting form factors were submitted to GAJOE for ensemble selection. GAJOE employs a genetic algorithm to iteratively select sub-ensembles from the initial pool whose average scattering best fits the experimental SAXS profile. Standard parameters were used, including a pool size of 9120 conformers, a maximum ensemble size of 20 conformers and a minimum of 5, with curve repetition enabled, and 100 generations of selection. Fitness was evaluated by minimizing the χ^2^ discrepancy between experimental and calculated scattering profiles, and convergence criteria were applied according to the ATSAS documentation (Manalastras-Cantos et al., 2021). Following optimization, global parameters including the radius of gyration (Rg), maximum particle dimension (Dmax), and Cα-Cα end to end distances were calculated for both the initial pool and the ensembles selected by GAJOE. Distributions of these parameters were used to quantitatively describe the conformational variability of the system, and the representative models that constitute the ensemble have been extracted for visualisation purposes. All calculations were performed using ATSAS package and default EOM pipeline.

## Results

### SAXS characterization of the two individual components

We started studying the structural behaviour of the protein by SAXS in the hope to capture new information on its dynamics. The scattering profile of the isolated double-stranded ARG-1A DNA aptamer was consistent with that of a compact duplex (**Figure 1A, Table 1)**, with a Guinier radius of 12.96 ± 0.05 Å and a maximum dimension (Dmax) of 46 Å, although an accurate determination of the Rg was complicated by a non-negligible Structure Factor contribution (**Figure 1B**), owing to the strongly repulsive interactions between negatively charged DNA duplexes at this concentration. The aptamer’s Molecular Weight (MW) was estimated as 12.1 kDa using the Volume of Correlation (Vc) method in RAW (Rambo & Tainer 2013), which is close to the theoretical 14.7 kDa, considering the difficulties in the determination of Vc for such a small particle. Inverse Fourier Transform of the scattering curve yielded a bell-shaped P(r) with a sharp decay (**Figure 1D**), confirming the compactness of the DNA molecule. No evidence of aggregation or structural heterogeneity was observed at this level.

**Table 1.**
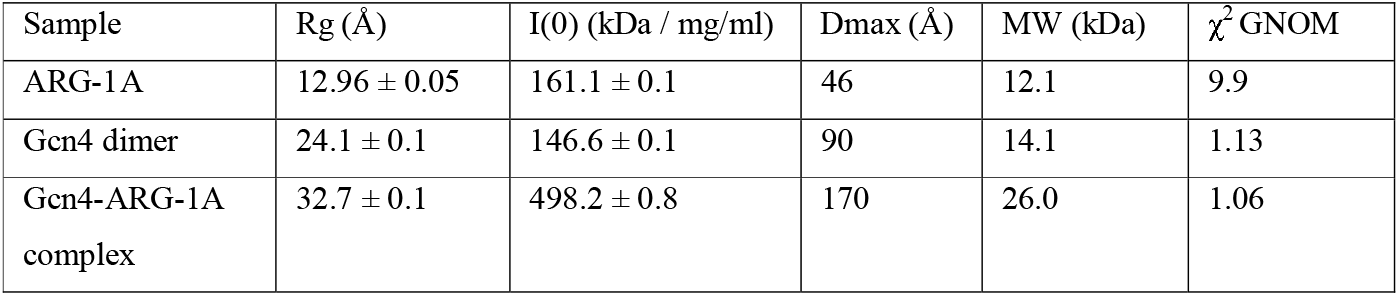
Summary of the structural parameters including Rg, Dmax, I(0), molecular weight estimates using the volume of correlation method, and χ^2^ values from GNOM fits (Svergun, 1992).

| Sample | Rg (Å) | I(0) (kDa / mg/ml) | Dmax (Å) | MW (kDa) | $\chi^2$ GNOM |
| --- | --- | --- | --- | --- | --- |
| ARG-1A | 12.96 ± 0.05 | 161.1 ± 0.1 | 46 | 12.1 | 9.9 |
| Gcn4 dimer | 24.1 ± 0.1 | 146.6 ± 0.1 | 90 | 14.1 | 1.13 |
| Gcn4-ARG-1A complex | 32.7 ± 0.1 | 498.2 ± 0.8 | 170 | 26.0 | 1.06 |

**Figure 1.**
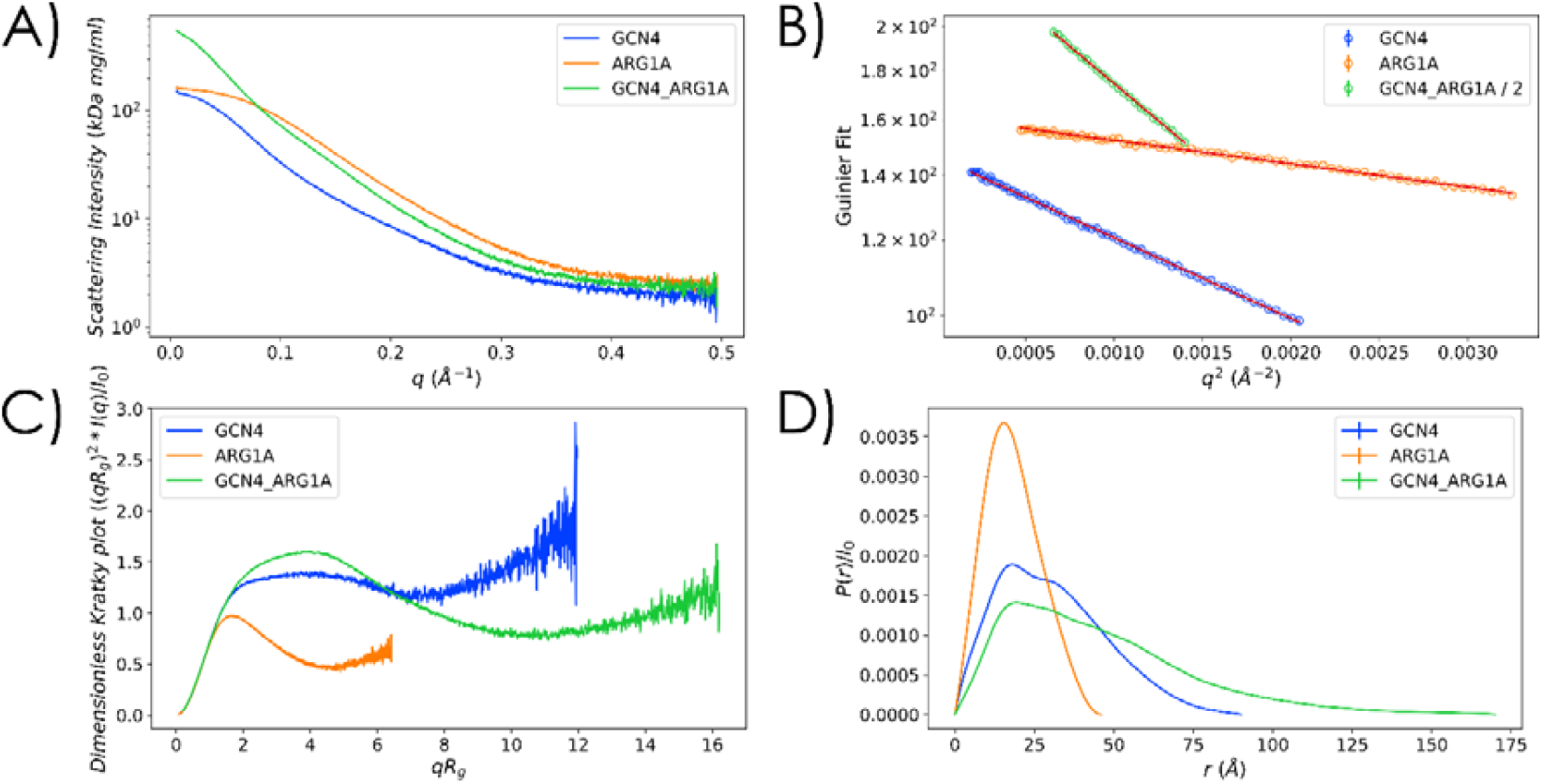
SAXS analysis of Gcn4 and the Gcn4-ARG1A DNA complex. **A**) Experimental scattering curves for the ARG-1A DNA aptamer (orange), the Gcn4 leucine zipper dimer (blue), and the Gcn4-DNA complex (green). **B**) Guinier fit plots. Note that the plot for the complex (green) has been divided by 2 to shift it on the log scale, for clarity. Error bars are within the symbol size. **C**) Dimensionless Kratky plots highlighting compactness of the DNA aptamer, partial disorder in Gcn4, and reduced flexibility in the complex. **D**) Pair distance distribution functions P(r) for each sample.

The scattering profile of the Gcn4 bZIP dimer indicated a substantially larger particle than the DNA aptamer (**Table 1**), with a Guinier radius of 24.1 ± 0.1 Å, and a Dmax of 90 Å. The molecular weight determined from the volume of correlation was 14.1 kDa, in good agreement with the theoretical dimer value of 13.7 kDa. The Kratky plot displayed a plateau at intermediate q-values, highlighting the decay in the Porod region of the scattering curve, characteristic of cylinder-like structures such as the coiled-coil motif (**Figure 1C**). The uptick at higher q-values indicates the presence of conformational disorder, which correlates with the expected disordered conformation of the basic region of Gcn4 in the absence of DNA. The P(r) shows a peak and an overlapping shoulder of slightly lower height consistent with the stable dimeric structure of Gcn4 bZIP, followed by a slowly decaying tail arising from the disordered basic region (**Figure 1D**).

We also performed a flexible refinement of Gcn4 bZIP, obtained using the software SREFLEX (Panjkovich and Svergun, 2016) on the SAXS data of the protein alone (**Figure 2A and S4**). SREFLEX from the ATSAS suite refines high-resolution macromolecular models by exploring the conformational space and improving their fit to the experimental SAXS data. It is based on a hybrid approach that combines normal mode analysis to identify potential conformational changes and then partitions the structure into pseudo-domains to analyze large and small movements, refining the structure to better match the experimental scattering profile. The coordinates of the protein were extracted from the PDB structure (1YSA), disregarding the DNA coordinates. SREFLEX was run with default parameters, in both constrained and unconstrained modes, and the best-scoring conformer from the unconstrained run was selected, after assessing the absence of breaks and steric clashes in the model. The resulting model displayed a remarkably good fit to the data (χ^2^ = 1.58, to be compared with a 21.21 value obtained for the starting 1YSA model) (**Figure S4**).

**Figure 2.**
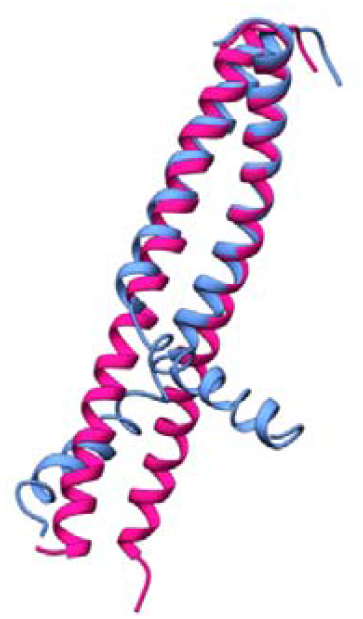
Superposition of the structure after treatment witb SREFLEX and the X-ray sf GCN4 bZIP. The kink and the deviation from a straight helix occurs at the interface between the DNA binding domain and the leucine zipper.

### Interaction between Gcn4 bZIP and ARG-1A leads to phase separation

We then set to study the properties of the bZIP-DNA complex using the ARG-1A DNA aptamer, which has high affinity and specificity for Gcn4 (Coey and Clark, 2022). Unexpectedly, when Gcn4 was mixed with the ARG-1A aptamer, the sample became immediately cloudy (**Figure 3, left panel**). It should be noted that the bZIP of Gcn4 has an equilibrium dissociation constant (*K*_d_) with its cognate DNA of 8 nM (Zitzewitz et al., 1995). Dimer formation is therefore expected to be strongly stabilized under the adopted experimental conditions. To understand the effect of pH, we repeated the mixing both at pH 7.4 and at pH 5.7, which are the conditions indicated in the paper by Ellenberger et al. (1992). These authors crystallised a DNA complex of this domain at the same protein concentration and in the same buffer (MES), although using a slightly different DNA aptamer. We observed similar effects at both pH values (5.7 and 7.4), with the formation of a highly cloudy solution indicating phase separation.

**Figure 3.**
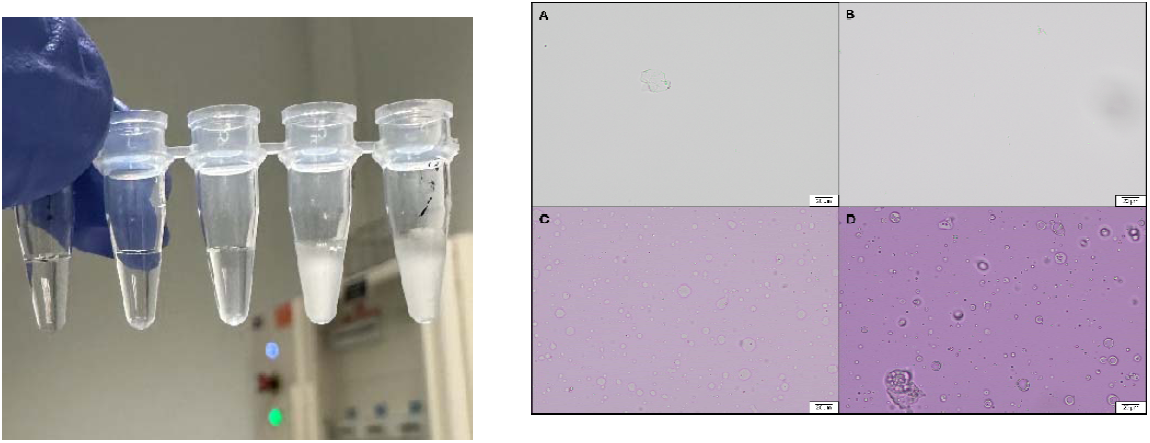
LLPS of the complex Gcn4 bZIP-ARG-1A. Left panel: The two rightmost samples contain mixture of Gcn4 with the aptamer. The samples became cloudy immediately after mixing. This occurred both at pH 7.4 and pH 5.7 that are the same conditions used in the paper by Ellenberger et al (1992). The authors crystallised the complex at the same concentrations and in this same buffer used in our measurements (MES). **Right panel**: Optical microscopy carried out on the isolated bZIP (a), the isolated ARG-1A (b), and the complex (c and d are two different regions of the same sample).

To characterize the solution and understand what sort of phase separation we were observing, we used optical microscopy. We measured individually ARG-1A and Gcn4 bZIP alone, and the complex (**Figure 3, right panel**). Nothing noticeable was observed for the two individual components except for a few dust particles. For the complex, we observed a change in turbidity with the appearance of a cloudy white material. During the measurement, using a 20× magnification, we observed many droplets and circular particles ranging in size from 2 µm to 15 µm. The smallest particles were in motion, while the largest ones were stable.

These results demonstrated that the bZIP domain has a strong tendency to phase separate and is sufficient to induce the transition. This is in contrast with previous reports in which the authors saw LLPS only in samples of Gcn4 constructs containing the transcription activator domain (Bremer et al., 2025). However, what we observed makes sense in terms of the physico-chemical characteristics of the protein.

### SAXS allows us to probe the complex also in a phase-separated sample

The SAXS curve for the Gcn4-ARG-1A complex was not significantly impacted by phase separation, except at very low q values (**Figure 1, Table 1**). This is because the radius of the droplets is in the 1-10 µm range, and thus no significant contribution from their form factor is expected in most of the q range of the instrument. Moreover, the droplets are smaller than the beam size (200x200 µm on BM29), therefore our data must reflect the average of the complex conformation in both the dilute and the droplet phases. Guinier analysis revealed an Rg of 32.7 ± 0.1 Å, while the maximum dimension was 170 Å, that is nearly twice that of the protein alone. The molecular weight estimated by the Vc method was 26.0 kDa, that is close to the theoretical 28.4 kDa for a Gcn4 dimer bound to a DNA duplex. However, this value should be considered only a qualitative estimate: while the Vc method can accurately estimate the electron density values for proteins or nucleic acids separately, it cannot produce accurate estimates when they are present in a mixture (Rambo & Tainer 2013).

The Kratky plot relative to the complex appreciably differed from that of the isolated protein as it showed a more rounded curve as compared to free Gcn4, owing to the more complex shape of the resulting structure (**Figure 1C**), while the more pronounced decay at higher q values suggested reduced flexibility upon DNA binding. The P(r) distribution resembled that of the protein alone at low r values, suggesting that the dimeric structure of the protein is, as expected, preserved, and it displays additional density at higher r values, indicating strong binding of the aptamer to the protein in agreement with what is experimentally known (**Figure 1D**).

### Simulating the dynamics of Gcn4 bZIP

The EOM strategy was applied to investigate the conformational properties of the Gcn4 bZIP–ARG-1A complex in solution (**Figure 4**). This widely adopted method is used to determine the structure of flexible proteins and other molecules that are not captured by a single static model (Bernardó et al., 2007; Tria et al., 2015). In the method, a large pool of potential conformations is generated by random sampling of the conformational space of a bead model, from which a subset of conformations is selected by a genetic algorithm. The goal is to find an ensemble of conformations whose combined theoretical SAXS scattering profile best matches the experimental data. In our case, the initial pool of conformations to test against experimental restraints was generated by a 100 ns all-atom molecular dynamics simulation trajectory run on the Gcn4 bZIP complex with ARG-1A, from which we extracted 9120 frames. It is worth of notice that the use of an all-atom representation rather than a coarse-grained model, as usually used by other authors for EOM calculations, gave us the advantage of sampling the conformational space in a physically more meaningful way, taking into account all the interactions that govern the binding process.

**Figure 4.**
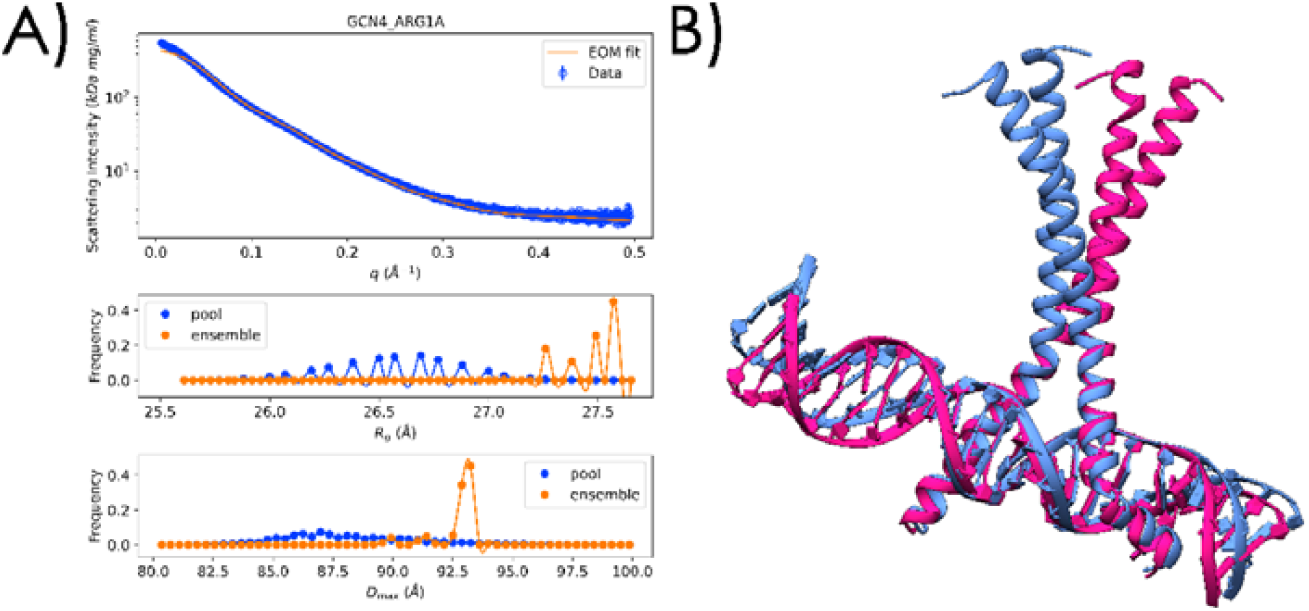
Ensemble Optimization Method (EOM) analysis of the Gcn4-ARG-1A complex. **A**) Top panel: Experimental SAXS curve (blue) and best EOM fit (orange). Middle panel: Distributions of radius of gyration (Rg) for the initial pool (blue) and optimized ensembles (orange). Bottom panel: Distribution of the maximum particle dimension (Dmax) for the initial pool (blue) and optimized ensembles (orange). **B**) Ribbon representation of the two main conformations identified by EOM. The magenta conformation has a weight of 90% and resembles the crystal structure, while the blue structure has a weight of 10% and presents a helical bend.

The simulations showed that the protein tends to breathe despite retaining its dimeric structure. During the trajectory, the helical structure is also preserved as indicated by the analysis of the secondary structure (Kabsch and Sander, 1983) and the root mean square fluctuations (RMSF) (**Figure S5**). However, occasionally the helices bend at the level of the interface between the DNA binding region and the leucine zippers (around residue 234), and this will be further explored in the following sections.

### Gcn4 bZIP exists in two different conformations

The experimental SAXS profile was excellently reproduced by the EOM fit, indicating that the selected ensembles provide a satisfactory description of the scattering data (**Figure 4A**). We noticed however a non-negligible discrepancy of the fit at low q and a fit χ^2^ of 214, when calculated manually. These deviations must reflect the phase separation process occurred during the component mixing.

The ensemble selected by the genetic algorithm consisted of a total of 10 curves, one of which repeated 9 times. This indicates the presence of two main conformations, with a relative population of 9:1. The dominant conformation closely resembles the crystal structure (1YSA), in which the protein helices are straight (**Figure 4B**). The minor conformation presents a kink at the end of the coiled-coil region which has not been previously described. The global parameter distributions (Rg and Dmax) of the pool resemble bell-shaped curves, indicating a good sampling of the conformational landscape of the complex (**Figure 4A**). The selected ensemble distributions of both parameters are much narrower, indicating that the solution structure of the complex is well described only by a restricted subset of the simulation frames. In particular, the average Rg of the ensemble is 27.5 Å, that is marginally bigger than the pool value of 26.6 Å. The ensemble average Dmax is 92.9 Å, versus the 88.4 Å value of the starting pool. This suggests that the experimental data are well modelled by selecting more elongated conformations from the pool, as it could be observed by the significantly higher weight of the “straight” conformation.

To estimate the system flexibility, we used the Rflex metric as defined in Burger et al. (2016). Rflex is defined as the ratio of the standard deviations of a selected ensemble and that of a pool. A value of Rflex of 100% indicates a fully flexible system while Rflex of 0% indicates a fully rigid system. We calculated an Rflex of 61.1% for the random pool, whereas the selected ensembles exhibited a significantly reduced value of 31.8%, consistent with a restricted conformational landscape (**Table 2**).

**Table 2.**
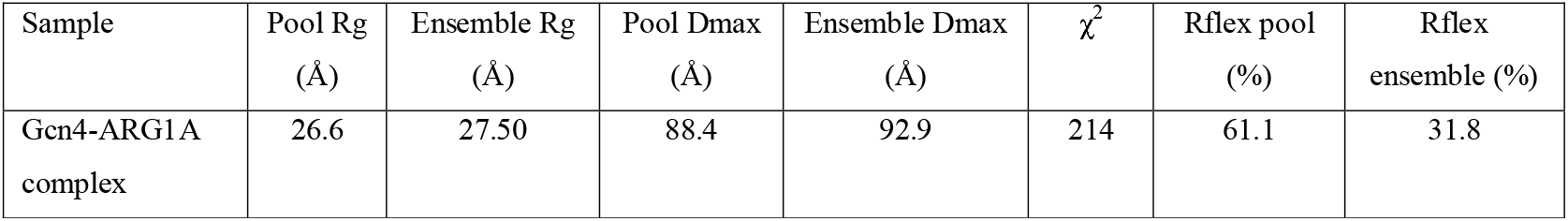
Summary of structural parameters derived from EOM, including Rg, Dmax, and flexibility indices (Rflex).

| Sample | Pool Rg<br>(Å) | Ensemble Rg<br>(Å) | Pool Dmax<br>(Å) | Ensemble Dmax<br>(Å) | $\chi^2$ | Rflex pool<br>(%) | Rflex<br>ensemble (%) |
| --- | --- | --- | --- | --- | --- | --- | --- |
| Gcn4-ARG1A<br>complex | 26.6 | 27.50 | 88.4 | 92.9 | 214 | 61.1 | 31.8 |

### Comparison of the experimental evidence with simulations and independent validation

The kinking tendency of the coiled-coil motif with respect of the basic region highlighted by EOM was also noticed by a detailed inspection of the molecular dynamics trajectory. To quantify the helix bending, we plotted the angles between the axes of the N- and C-terminal regions of the helices along the trajectory, as defined in the methods section (**Figure 5**). Values close to 180° indicate a straight helix, while lower values indicate the presence of a kink. The plot that reports the values of the dihedral angles N1C1-N3C3 versus N2C2-N4C4 is rather dispersed, suggesting appreciable dynamics, with a main, dense cluster around values (155º, 170º), and a less dense one around (135°, 150°). The values of these angles for the kinked conformation selected by EOM fall into the latter cluster, reasonably close to its centre. Interestingly, the values for both the straight EOM conformation and the crystal structure are located at the very edge of the main cluster, indicating that the vast majority of the conformations visited by the dimer in the complex according to the molecular dynamics correspond to a bent dimer.

**Figure 5.**
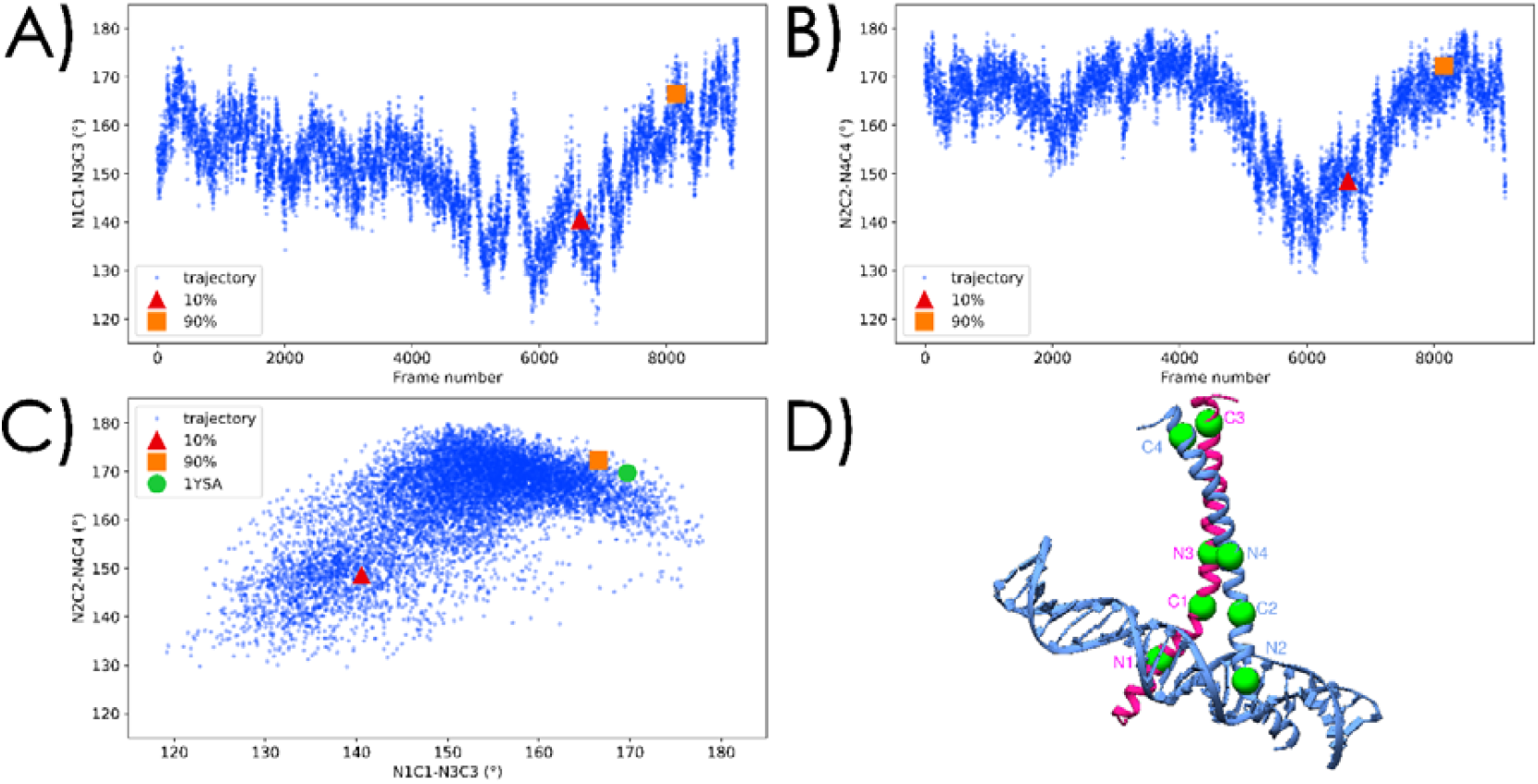
Analysis ot the interhelix angles of the monomer in the dimer angles along the trajectory. Variation of the N1C1-N3C3 (panel **A**) and N2C2-N4C4 (panel **B**) interhelical. The selected EOM structures are highlighted. **C**) Two-dimensional scatter plot of the helical axis angles N1C1-N3C3 and N2C2-N4C4. The values for the kinked conformation from EOM (red triangle), the straight conformation (orange square) and the crystal structure (green circle) are shown. **D**) The selected residues that define the angles between the two helices shown as green balls (see definition in the text).

To further validate the results, we used the DENSS (DENsity from Solution Scattering) software (Grant, 2018). This is an *ab initio* modelling method that reconstructs a 3D particle density map directly from scattering data, bypassing the need of pre-existing atomic models. This approach thus solves the inverse problem by iteratively using the experimental scattering intensity to refine a density map. The overlayed electron densities calculated with DENSS showed that the structure of the unbound protein is stable in the coiled-coil region, with its disordered DNA binding domains elongating away from each other due to their electrostatic repulsion (**Figure 6A**). Conversely, the electron density of the complex loosely follows the overall shape of the model (**Figure 6B**). The conformational change and the disorder-to-order transition in the basic region of the Gcn4 bZIP upon DNA binding is evident from this analysis. Interestingly, the residues where the kink is present in the bound state are those where the disordered region starts in the unbound state and thus in the interface between the DNA binding domain and the leucine zipper. It is also clear that both the dominant straight and the kinked conformations of the Gcn4 bZIP in the complex can comfortably fit the calculated density map.

**Figure 6.**
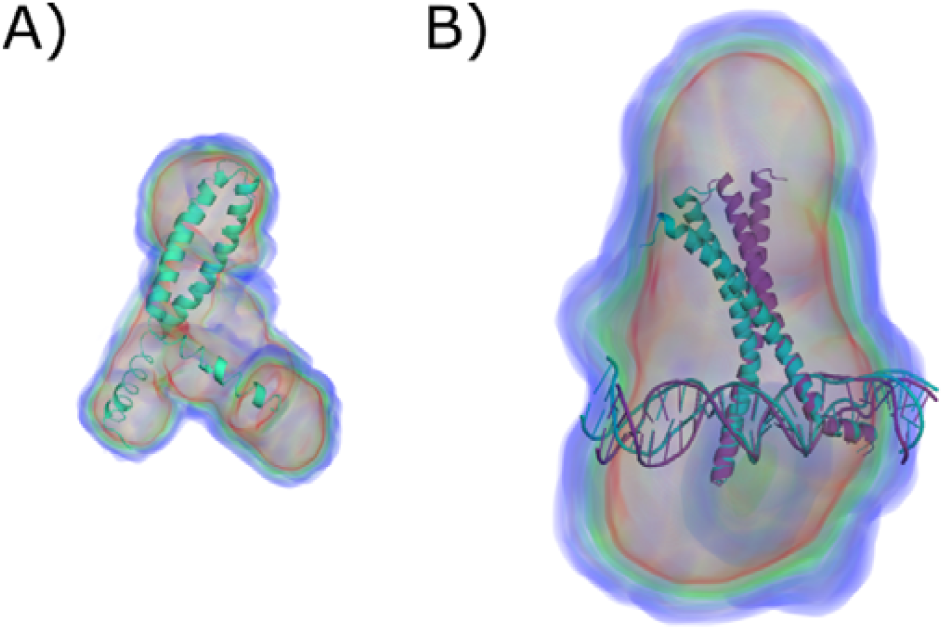
Fitting of the Gcn4 bZIP domain into the SAXS envelope. **A**) Flexible refinement of the Gcn4 dimer using SREFLEX, overlayed with the electron density calculated with DENSS. **B**) representative conformations from the final EOM ensemble, illustrating straight (magenta) and kinked (cyan) states of the Gcn4-ARG-1A complex, overlayed with the electron density of the complex calculated using DENSS from the experimental data.

Together, these results provide a novel and more accurate description of the conformational transitions and the dynamics undergoing during complex formation and DNA recognition of Gcn4 bZIP.

## Discussion

In the present study, we explored the conformational properties in solution of the bZIP domain of Gcn4, a well-known transcriptional activator that has been studied for more than three decades for several completely different purposes. Initially interesting because of the presence of the first coiled-coil domain in a protein other than a muscle-related one, GCN4 then became a powerful tool in novel protein design and in many applications in which a dimeric structure was needed (Weissenhorn et al., 1997; Wolfe et al., 2003; Mason and Arndt., 2004; Deiss et al., 2014). Gcn4 is also an excellent example of how disordered or partially disordered transcription factor effector domains can be important in transcriptional regulation (Datta et al., 2025). More recently, Gcn4 has attracted renewed interest to shed light onto the debate about the role of phase separation in transcription (Bremer et al., 2025). It is in fact currently heatedly debated whether LLPS is an event necessary for transcription or occurs only *in vitro* or under specific circumstances. Bremer et al. (2025) used Gcn4 to assess the possibility of a mechanism mediated by soluble complexes or by condensates and found that the ability of Gcn4 to form soluble complexes with its coactivator Med15 correlates with its propensity to recruit this protein into condensates, giving some support the importance of LLPS in transcription.

We originally started to work on Gcn4 within a project designed for a completely different purpose: we were aiming at studying the interactions of proteins with DNA plasmids with different supercoiling. To do this, we first characterized as a reference the structure of Gcn4 bZIP in the presence and in the absence of a small DNA aptamer spanning the Gcn4 recognition sequence not expecting to find anything interesting beyond what had already been extensively described. We observed instead a very interesting and rather unexpected phenomenon: when we mixed DNA with the protein, we observed phase separation. This was unexpected because Bremer et al. (2025) had excluded the possibility of such transition for the isolated DNA binding bZIP motif which, in their studies, was used as a negative control. They attributed the LLPS properties only to the transcription activator domain downstream to bZIP. They ascribed this behaviour to the intrinsically unfolded character of the transcription domain. Yet, our data show that also the bZIP domain undergoes the transition. this is interesting because it underlines an overall ability to phase separate throughout the whole sequence of Gcn4 that is independent of the presence of Med15. The differences between the two studies could stand in the length of the bZIP construct (we used 226-281 versus 222-281), in a slightly different DNA sequence embedding the same Gcn4 binding consensus sequence, and/or in slightly different experimental conditions. Showing that DNA binding is sufficient to trigger LLPS of the bZIP reframes how we think about charge patterning, DNA–protein coacervation, and the minimal requirements for condensate formation in transcription factors.

It is certainly tempting to suggest why we observe this phase transition: it is well known that a distinctive form of electrostatically driven LLPS known as complex coacervation, occurs when two or more oppositely charged macromolecules interact in aqueous solution because they neutralise each other. As a consequence, the system spontaneously divides into two liquid phases: a dense, polymer-rich phase known as the coacervate, and a dilute, polymer-poor supernatant. This transition does not produce a solid or gel, but rather a set of droplets that retain liquid-like properties, including internal mobility, the ability to fuse, and the rapid exchange of molecules with the surrounding medium, that is exactly what we observed. In our case, we were mixing the negatively charged DNA with a positively charged protein (with an estimated pI of 10.95). These considerations remind us of an important message: intrinsically disordered proteins and protein domains do have a high tendency to phase separate but disorder is neither necessary nor sufficient to have to observe LLPS. The transition may occur also with protein regions that are mostly folded, as it is the case for the bZIP, because the forces that promote the transition are quite diversified (Aprile et al., 2021).

We then used a combination of SAXS and all-atoms molecular dynamics to get structural information. While low resolution as compared to crystallography, nuclear magnetic resonance or cryo-electron microscopy, SAXS is a technique particularly powerful to describe the overall picture of a complex and to highlight conformational changes. In our case, we found that SAXS could not only allow us to describe the known protein conformation but also obtain direct and quantitative information on the structure of the protein in- and outside molecular condensates. This is because in a SAXS measurement, all the different conformations of the molecules that are present in the illuminated volume equally contribute to the measured signal, giving a snapshot of the whole conformational ensemble that these molecules can explore. The use of all-atoms simulation to interpret the SAXS data offered on the other hand the novelty of using a full atomic description of the system. This is at variance with the vast majority of similar studies that rely on a coarse-grain representation of the molecular dynamics (see for instance Martin et al., 2022). Our, together with Hernandez et al. (2025), is one of the very few examples of a EOM study based on an all-atoms representation.

Our data of the isolated bZIP domain conclusively reflect its disordered and dynamical properties: the data indicate the presence of local conformational disorder within a molecule that forms a stable dimeric structure. These fundings are in full agreement with previous NMR studies (O’Shea et al., 1991; Saudek et al., 1991; Ellenberger et al. 1992), demonstrating once again how powerful SAXS can be also to study intrinsically disordered proteins (Uversky, 2025).

The analysis of the complex is even more interesting. While the crystal structure of the bZIP-DNA complex (Ellenberg et al., 1992) gives us a static picture of two uninterrupted helices grasping in the middle the DNA, our data provide a more nuanced picture in which we have indications for the first time of the presence of two populations, the most represented one is in agreement with the crystal structure, the second, much less represented, has the helices slightly bend at the level of the transition from the basic region to the leucine zipper motifs. EOM reveals that this conformation accounts for only 10% of its solution conformational ensemble within our experimental conditions, while the remaining 90% is represented by straight, extended structures. These two populations could of course reflect the presence of a mixture of a mixture of two conformations with similar dynamics both in the diluted phase and in the condensate droplets. However, the close similarity of the straight conformation with the crystal structure could suggest an interesting alternative explanation: the straight dimer could be the dominant conformation in the droplet phase, where the complexes would be closely packed because of confinement, as suggested by the conceptual similarity of this situation with that in the crystal. The kinked conformation could dominate instead in the dilute phase, in agreement with our molecular dynamics simulations that indicate a strong tendency of the helices to bend in an aqueous environment. It may be noticeable that the kink appears exactly at the order-disorder boundary, making it mechanistically plausible and potentially relevant for DNA binding and flexibility. Against this explanation, it could be objected that there is a relatively modest difference in the population ratio (9:1) of the two form while one could expect a much bigger ratio in favour of the protein in the droplet since its concentration should be intrinsically much higher. This apparent discrepancy could easily be rationalized by contrast effects, as complexes in the dilute phase experience a much higher scattering contrast relative to the solvent than complexes embedded within the protein-rich droplet phase, where the surrounding medium has a similar electron density to the protein itself.

In conclusion, our results provide new and exciting insights into the conformational space of Gcn4 in isolation and in a complex with DNA and represent a further example of how powerful SAXS can be in accurately and quantitatively describing also minute conformational differences and minor populations also when the sample is phase separated. Rather than treating LLPS as an experimental nuisance, we exploited the ability of SAXS to average over dilute and droplet phases to extract new information on all the structural populations present in solution and connected these populations to distinct physical environments (crowded/droplet-like *vs* dilute phase), something crystallography and even NMR cannot easily access. Our data thus help to understand the mechanism of the LLPS transition of Gcn4 and open a new chapter into the possibility to characterize structurally proteins in coacervate droplets.

## Acknowledgments

We sincerely thank Arthur Palmer III for the generous gift of the GCN4 bZIP plasmid. We are strongly indebted with Jonathan Leon Moro, Anna Valenti and Mikael Widersten for discussions and suggestions. The work was supported by the European Innovation Council Pathfinder Open project “iSenseDNA” cod. 101046920. The computations and their analysis were enabled by resources provided by the National Academic Infrastructure for Supercomputing in Sweden (NAISS), partially funded by the Swedish Research Council through grant agreement no. 2022-06725. We thank ESRF for access to the BM29 beamline.

