## Supplementary material for "DNA binding drives phase separation of the Gcn4 bZIP domain and reveals its conformational ensemble in the diluted and condensate phases": Suppl. Figures S1-S5

**Supplementary Materials**

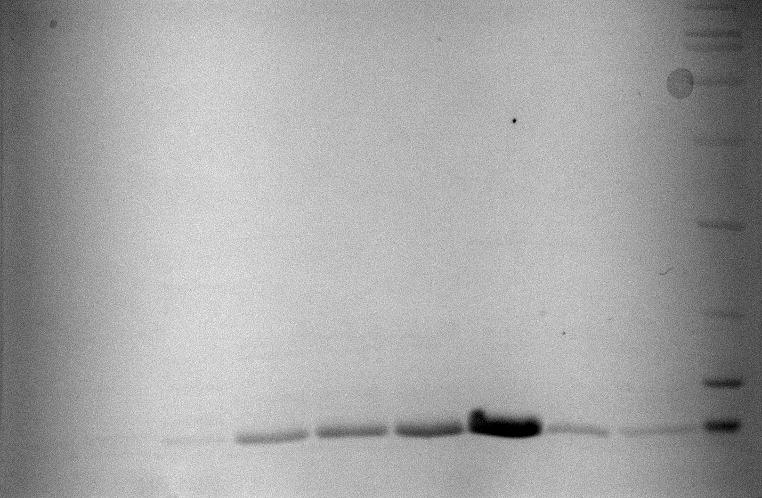

**Figure S1. SDS-PAGE of Ion Exchange Chromatography fractions after purification.** The electrophoretic profile showed a predominant band at the expected molecular weight, with no detectable higher molecular weight contaminants, confirming the efficiency of the purification procedure.

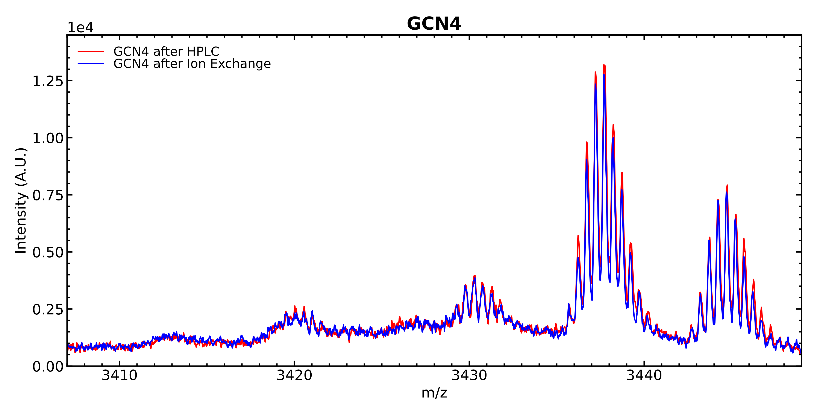

**Figure S2 – MALDI-TOF Spectra for doubly ionized Gcn4 after the two independent purification protocols with (blue) or without (red) an extra step of purification.** The spectra confirmed a mass consistent with the theoretical monomeric species and indicated a purity above 98%. They also confirmed that the ion-exchange purification alone is sufficient to obtain highly pure material.

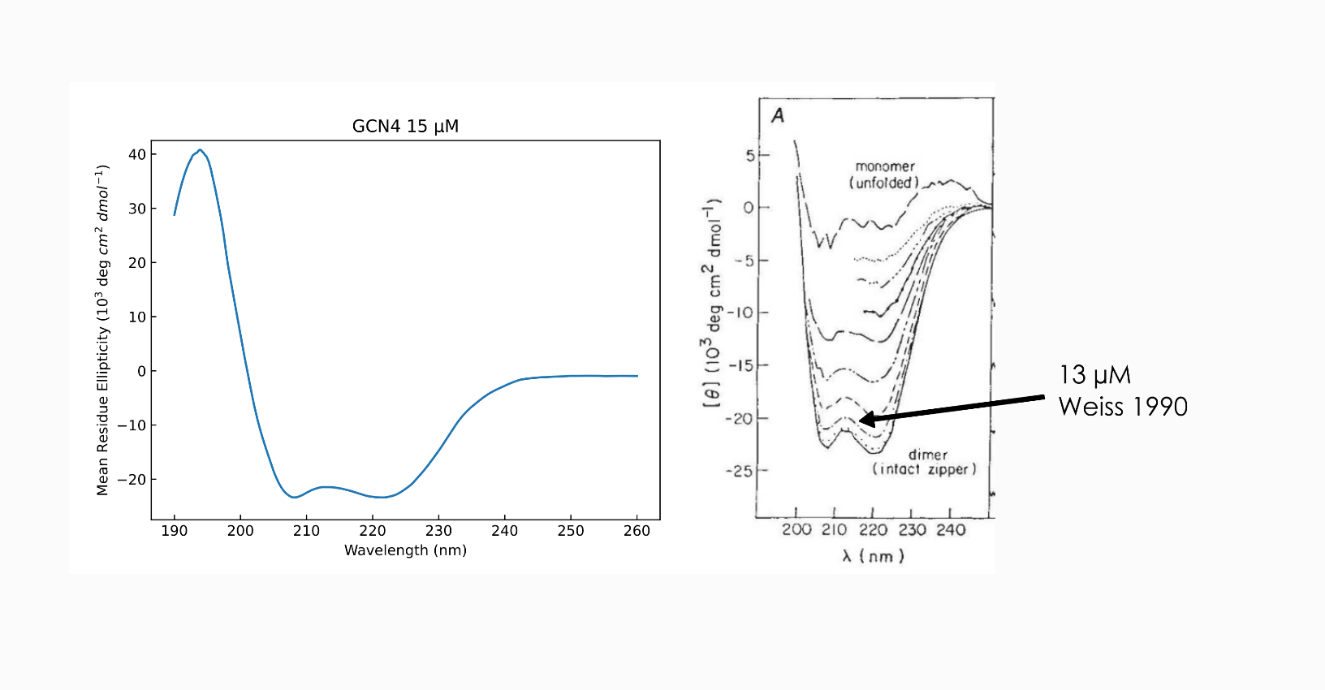

**Figure S3 – Normalised far-UV CD spectrum** of Gcn4 bZIP. The experiment was carried out on a JASCO-1100 spectropolarimeter equipped with the Peltier temperature control system measured in 1 mm pathlength quartz cuvettes (type S3/Q/1; Starna Scientific). Protein concentration was 15 µM and the buffer composition was 25 mM HEPES, 50 mM NaCl, pH 7.5. Spectra were recorded with a step size of 0.1 nm, a bandwidth of 1 nm, and a scanning speed of 50 nm/min. Ten consecutive scans were collected, averaged, and baseline corrected against buffer spectra. The protein displayed the expected typical α-helical signature with two minima at 208 and 222 nm, in agreement with previous reports of Gcn4 (Ellenberger et al., 1992). The normalized spectrum corresponds to a ca. 70% helical content according to the literature (O’Shea et al., 1989).

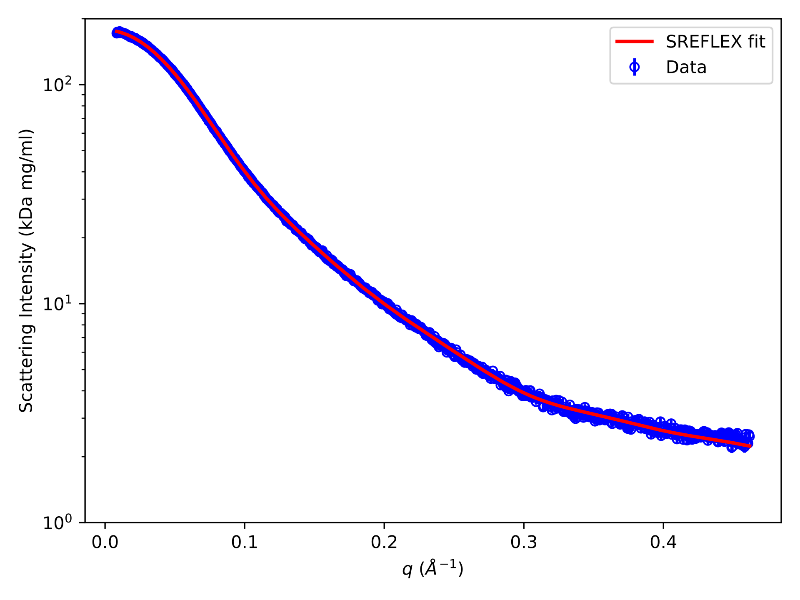

**Figure S4** – **SREFLEX analysis.** Experimental SAXS curve for Gcn4 bZIP alone (blue line) and corresponding fit from the best-scoring SREFLEX model (red line).

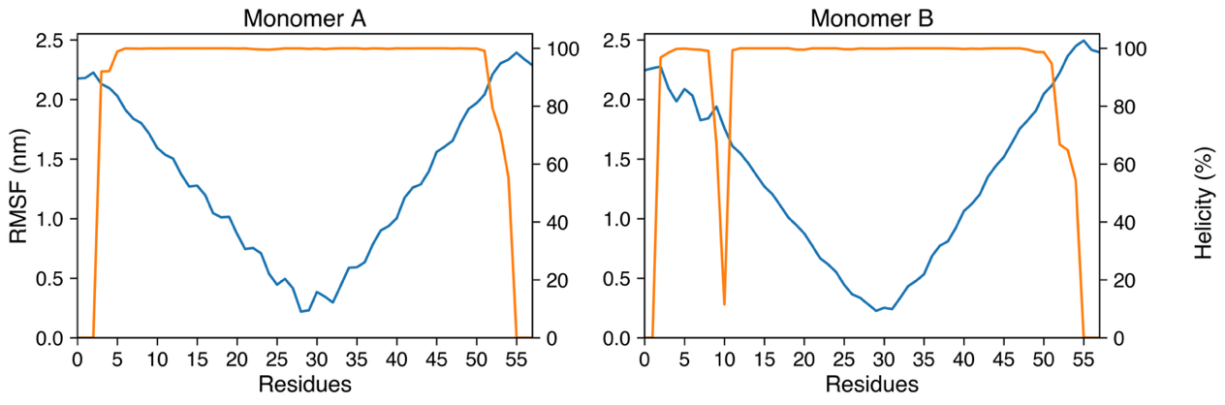

**Figure S5.** Analysis of the secondary structure of Gcn4 bZIP along the trajectory. **Top panel.** Per-residue Cα RMSF (nm) and per-residue helicity (%) of Gcn4 computed over the trajectory for **Monomer A** and **Monomer B**. RMSF reports backbone flexibility, while helicity indicates the fraction of frames assigned to α-helix by DSSP.

Ellenberger TE, Brandl CJ, Struhl K, Harrison SC. Cell. 1992; 71:1223–1237. doi: 10.1016/s0092-8674(05)80070-4.

O’Shea EK, Rutkowski R, Kim PS (1989) Evidence that the leucine zipper is a coiled coil. Science 243:538–542.
